# CD5L associates with IgM via the J chain

**DOI:** 10.1101/2023.12.18.572142

**Authors:** Yuxin Wang, Chen Su, Chengong Ji, Junyu Xiao

## Abstract

CD5 antigen-like (CD5L), also known as Spα or AIM (Apoptosis inhibitor of macrophage), emerges as an integral component of serum immunoglobulin M (IgM). However, the molecular mechanism underlying the interaction between IgM and CD5L has remained elusive. In this study, we present a cryo-electron microscopy structure of the IgM pentamer core in complex with CD5L. Our findings reveal that CD5L binds to the joining chain (J chain) in a Ca^2+^-dependent manner and further links to IgM via a disulfide bond. We further show that CD5L reduces IgM binding to the mucosal transport receptor pIgR, but does not impact the binding of the IgM-specific receptor FcμR. Additionally, CD5L does not affect IgM-mediated complement activation. These results offer new insights into our understanding of IgM and shed light on the functions of the IgM–CD5L complex in the immune system.

---

Immunoglobulin M (IgM) plays a critical role in both humoral and mucosal immunity. In serum, IgM is primarily found in its pentameric form, consisting of five IgM monomers linked together by the joining chain (J chain). The J chain is also essential for the transportation of the IgM pentamer to the mucosal surface, and achieves this by interacting with the polymeric immunoglobulin receptor (pIgR), a part of which becomes an integral subunit of secretory IgM known as the secretory component (SC).

The discovery of IgM can be traced back to 1937 when Heidelberger and Pedersen observed that horses immunized with pneumococcus polysaccharide produced antibodies with a remarkably large molecular weight^1^. Despite its long history, our understanding of IgM remains incomplete. The presence of an additional IgM-associated protein in human serum IgM, apart from the J chain, was initially identified by Tissot et al. using two-dimensional polyacrylamide gel electrophoresis in 1993–1994^2,3^. Gebe et al. subsequently cloned and characterized human Spα in 1997 as a macrophage-secreted protein^4^, which belongs to the Scavenger Receptor Cysteine-Rich (SRCR) superfamily. Miyazaki et al. identified mouse AIM (Apoptosis inhibitor of macrophage) in 1999, and found that it can promote the survival of thymocytes against apoptosis^5^. It was soon realized that AIM is the mouse homolog of human Spα^6^. In 2002, it was confirmed that human Spα/AIM, also known as CD5 antigen-like or CD5L, corresponds to the additional protein found in IgM^7^. Recently, Oskam et al. performed state-of-the-art mass spectrometry analyses and demonstrated that CD5L/Spα/AIM is an integral subunit of human circulatory IgM, leading to the redefinition of the predominant form of circulatory IgM as a J-chain-linked IgM pentamer plus one CD5L [(IgM)_5_:(J)_1_:(CD5L)_1_]^8^. For consistency with this study and in compliance with UniProtKB recommendation, we will primarily refer to CD5L/Spα/AIM as CD5L in this study.

In the past two decades, a wide range of functions of CD5L have been documented^9,10^. CD5L is believed to act as a pattern recognition receptor (PRR) that recognizes a broad range of microbial pathogens, including both bacterial and fungal cells^11-14^, as well as endogenous harmful substances such as cell debris^15-17^. In addition to its role in inhibiting the apoptosis of thymocytes, CD5L has been shown to suppress the apoptosis of T cells and natural killer T cells in mice^18^, as well as hepatocytes after hepatic ischaemia-reperfusion (I/R) injury^19^. CD5L also regulates lipid biosynthesis and the pathogenicity of T helper 17 cells^20^. Furthermore, CD5L has been implicated in diseases such as obesity, atherosclerosis, and IgA nephropathy^21-23^. Although the molecular mechanism of CD5L function remains elusive, it is clear that it plays important roles in immune homeostasis and diseases.

The IgM pentamer is believed to serve as a carrier for CD5L, shielding it from renal excretion. However, the molecular basis of the IgM–J–CD5L interaction remains unclear, and the regulatory role of CD5L in IgM function remains to be further investigated. In this study, we present a cryo-electron microscopy structure of the IgM-Fc pentamer core in complex with CD5L (Fcμ–J–CD5L), shedding light on the organizational principles of the IgM–CD5L complex. Additionally, our findings demonstrate that CD5L diminishes IgM binding to the mucosal transport receptor pIgR, while leaving the binding of the IgM-specific receptor FcμR unaffected. Furthermore, we show that CD5L has no impact on IgM-mediated complement activation. These insights provide a more comprehensive understanding of human IgM.

## Results

### CD5L binds to IgM in a Ca^2+^-dependent manner

The SRCR proteins are characterized by one or multiple repeats of highly conserved SRCR domains^24,25^. Structural analyses have shown that SRCR domains commonly feature a dual-cation-binding site that plays a crucial role in ligand binding^26^. To biochemically characterize the IgM–CD5L interaction and explore its dependence on divalent cations such as Ca^2+^, which is typically found in serum (at levels of 1.3–1.5 mM in healthy human blood), we generated CD5L, as well as Fcμ–J, the pentameric IgM core comprising an IgM-Fc pentamer and the J chain, using HEK293F cells (Extended Data Fig. 1a). Surface plasmon resonance (SPR) experiments demonstrated that CD5L binds to immobilized Fcμ–J with a Kd of 30 nM in the presence of Ca^2+^ (Fig. 1a). In contrast, no significant interaction was observed between CD5L and Fcμ–J in the presence of the chelating agent EDTA (ethylenediaminetetraacetic acid, Extended Data Fig. 1b). These findings support the strong interaction between Fcμ–J and CD5L, and suggest that, akin to other SRCR–ligand interactions, CD5L binds to IgM in a Ca^2+^-dependent manner.

**Figure 1.**
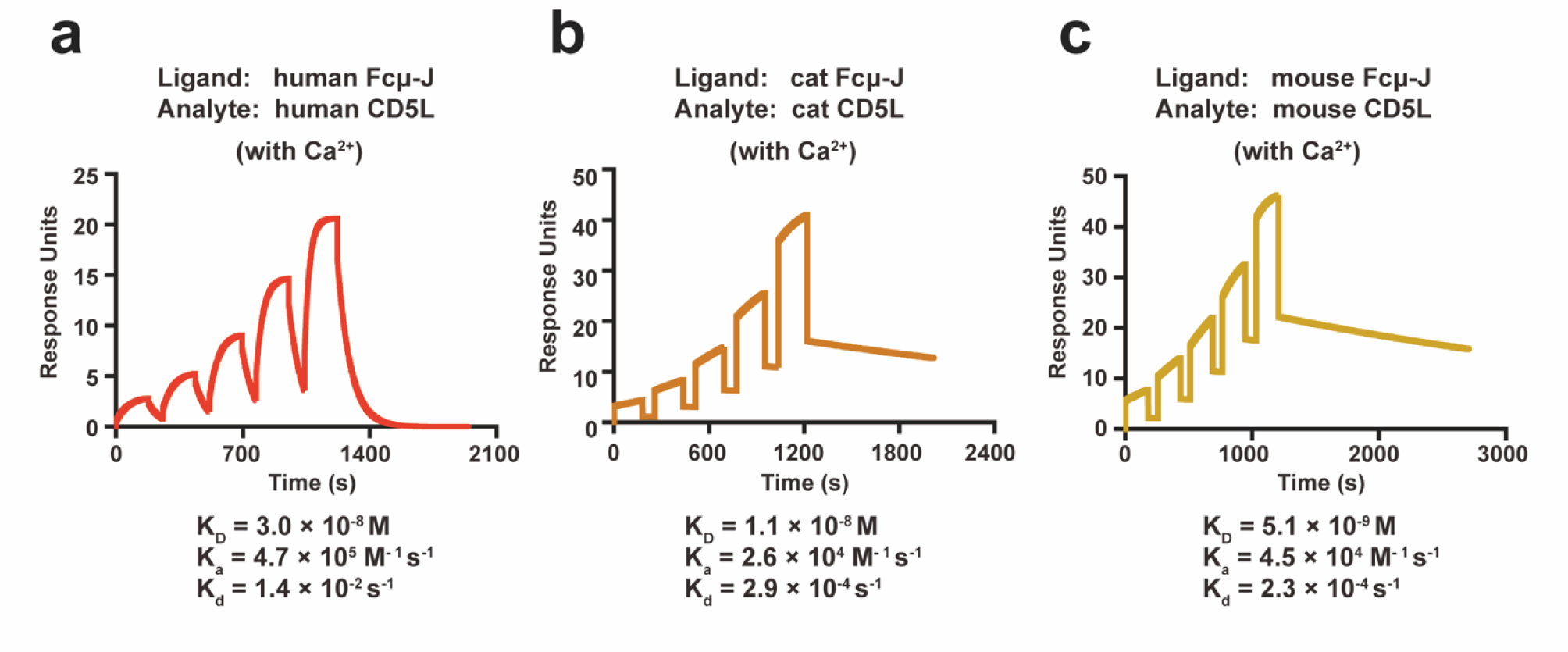
SPR analyses of the Fcμ–J–CD5L interactions. **a.** Human CD5L binds to immobilized human Fcμ–J with a Kd of 30 nM in the presence of Ca^2+^. The binding experiments were carried out using a single-cycle kinetics program. All SPR experiments in this paper have been repeated at least three times with similar results. **b.** Cat CD5L binds to immobilized cat Fcμ–J in the presence of Ca^2+^. **c.** Mouse CD5L binds to immobilized mouse Fcμ–J in the presence of Ca^2+^.

CD5L is highly conserved in mammals (Extended Data Fig. 2). Previous research has indicated that cat CD5L binds to cat IgM with a 1000-fold greater affinity compared to mouse CD5L and IgM^27^, and this strong IgM–CD5L binding in cats was suggested to contribute to their increased susceptibility to renal disease^10^. However, our measurements show that both cat and mouse CD5L exhibit similar affinities for their respective Fcμ–J, with Kd values of 11 nM and 5 nM, respectively (Fig. 1b, c).

### Cryo-EM structure of human Fcμ–J–CD5L

The core region of the IgM pentamer displays an asymmetrical arrangement that resembles a hexagon with a missing piece^28-32^. Negative-stain electron microscopy images indicate that CD5L binds to the gap of the IgM pentamer^28^. To reveal the molecular basis of IgM–CD5L interaction, we incubated human Fcμ–J complex and CD5L together to assemble the Fcμ–J–CD5L complex (Extended Data Fig. 1c), and then analyzed it using cryo-electron microscopy (cryo-EM). The overall structure was determined at a resolution of 3.4 Å (Fig. 2a, Extended Data Fig. 3, Extended Data Table 1), and revealed an asymmetrical arrangement of five Fcμ molecules with a gap occupied by the J chain and CD5L, in agreement with previous findings. The SRCR3 domain of CD5L binds to the center of Fcμ–J and interacts with several regions of the J chain. SRCR2 swings towards Fcμ5B, and presses the β5–β6 hairpin of the J chain onto Fcμ5B. When viewed from the side, the SRCR2–SRCR3 module was found to be non-uniformly distributed within the Fcμ–J plane, concentrating more on one side. This further accentuates the asymmetrical nature of the core region of IgM. Weak densities were observed for the SRCR1 domain of CD5L, as well as the Cμ2 domains of Fcμ, preventing clear visualization of these regions.

**Figure 2.**
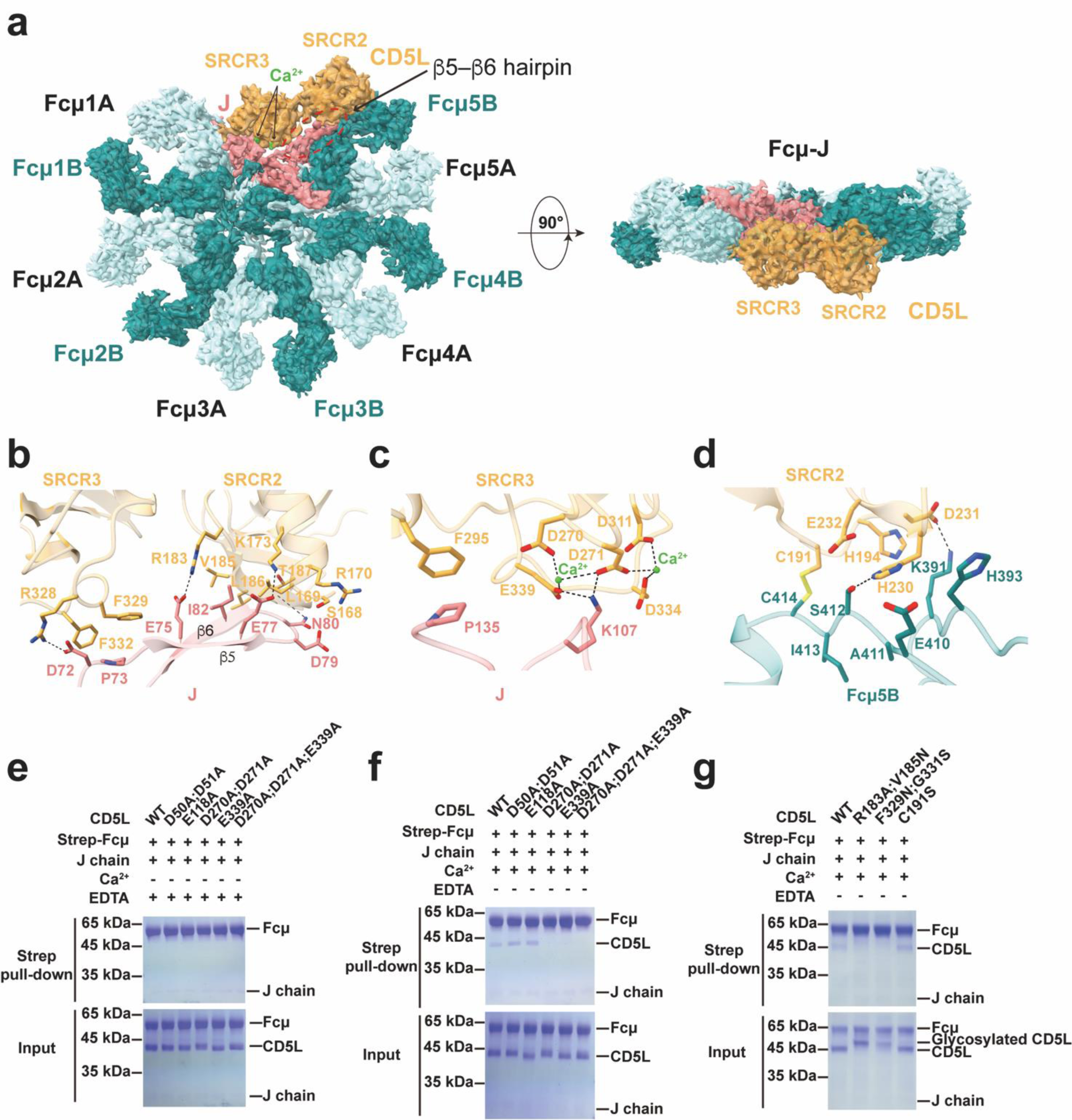
Cryo-EM structure of human Fcμ–J–CD5L complex. **a.** The cryo-EM structure of the human Fcμ–J–CD5L complex is shown in two views. Five Fcμ chains are shown in teal and five are shown in pale turquoise. J chain, CD5L and Ca^2+^ are shown in light coral, gold and green, respectively. Similar color schemes are used in all figures unless otherwise indicated. **b.** Interactions between the SRCR2–SRCR3 junction of CD5L and the β5–β6 hairpin of J chain. **c.** Interactions between DDE/D motif of the SRCR3 and J chain. There are two Ca^2+^ in the DDE motif and DDD motif of SRCR3. **d.** Interactions between the SRCR2 domain of CD5L and the Cμ3 of Fcμ5B. **e.** The interaction of CD5L and Fcμ-J complex is Ca^2+^-dependent. **f.** CD5L-SRCR3 mutants displayed abolished interaction with Fcμ–J. **g.** SRCR2 and SRCR3 mutants displayed reduced or abolished interaction with Fcμ–J.

Local refinement was performed to enhance the EM densities surrounding CD5L and the Fcμ–J–CD5L interface (Extended Data Fig. 3, Extended Data Table 1). Both the SRCR2 and SRCR3 domains of CD5L were found to be crucial in binding to the J chain, particularly the β5–β6 hairpin of the J chain (Fig. 2b). It is worth noting that this hairpin structure, observable in IgA structures^33-35^, was disordered in all previous IgM structures without CD5L. In SRCR2, K173 and R183 interact with E77_J_ and E75_J_ (J chain residues are denoted by subscripts), respectively; whereas V185–T187 pack with N80_J_–I82_J_. In SRCR3, R328 formed a salt bridge with D72_J_, while F329 and F332 attached to P73_J_.

Further interactions have been observed between SRCR3 and the C-terminal region of the J chain. Sequence analyses indicate that SRCR3 contains a complete dual-cation-binding site, while SRCR2 has a degenerate one^26^. Indeed, a cryo-EM density map reveals that a negatively-charged surface patch formed by D270, D271, D311, D334, and E339 in SRCR3 coordinates two Ca^2+^ ions (Extended Data Fig. 3g). A pocket between these two Ca^2+^-binding sites cradles Lys107_J_ (Fig. 2c). This observation aligns with the SPR analyses and demonstrates the crucial role of Ca^2+^ in facilitating the interaction between CD5L and Fcμ–J. In addition, F295 in SRCR3 interacts with P135_J_.

CD5L also directly interacts with Fcμ. In particular, a disulfide bond was observed to form between Cys191 in SRCR2 and Cys414 in the Cμ3 domain of Fcμ5B (Fig. 2d), consistent with previous analyses^8,28^. H230 and D231 from SRCR2 were also involved in making polar contacts with Fcμ5B. These interactions further contribute to the stabilization of the Fcμ–J–CD5L complex.

To validate the functional significance of these molecular interactions, we engineered CD5L mutants and conducted pull-down experiments. CD5L exclusively binds to Fcμ–J in the presence of Ca^2+^, and the interaction between CD5L and Fcμ–J is disrupted by EDTA (Fig. 2e, f). Furthermore, CD5L mutants carrying cation-binding mutations in SRCR3, such as D270A/D271A, E339A, and D270A/D271A/E339A, showed a complete loss of interaction with Fcμ–J. In contrast, mutations of the putative Ca^2+^-binding sites in SRCR1 (D50A/D51A and E118A) had no impact on the interaction between CD5L and Fcμ–J. The R183A/V185N and F329N/G331S mutations, engineered to introduce bulky N-linked glycans into amino acid positions 185 and 329, respectively, and thereby disrupt the interaction between CD5L and the J-chain β5–β6 hairpin (Fig. 2b), also substantially diminished the interaction between CD5L and Fcμ–J (Fig. 2g). Notably, mutation of C191 in CD5L did not significantly impact the binding between CD5L and Fcμ–J, indicating that the disulfide bond between this Cys and Fcμ5B is not the primary driving force for IgM– CD5L interaction. Instead, it likely functions to enhance the stability of IgM–CD5L in circulation.

### CD5L reduces IgM binding to pIgR but not to FcμR

The integration of CD5L as a subunit of IgM prompts an inquiry into its impact on IgM function. To address this, we started by investigating the influence of CD5L on the interaction between the IgM pentamer and the specific IgM Fc receptor, FcμR^36^. Surface plasmon resonance (SPR) experiments revealed that Fcμ– J–CD5L and Fcμ–J exhibit similar high-affinity interactions with immobilized FcμR, with apparent Kd values of 0.4 nM and 0.3 nM, respectively (Fig. 3a). This suggests that the presence of CD5L does not significantly affect the interaction between IgM and FcμR. Structural analyses further support this conclusion, indicating that CD5L is unlikely to impede the recruitment of FcμR by IgM (Fig. 3b). Similar conclusions were drawn from SPR experiments conducted in the reverse manner, where Fcμ–J–CD5L and Fcμ–J were immobilized on the SPR chip, albeit with lower apparent affinities (Kd values of 8.6 nM and 7.2 nM, Extended Data Fig. 4a). The observed affinity differences are likely due to avidity effects, as a single Fcμ–J can engage with up to four FcμR molecules on either side^37^. When FcμR is anchored to the SPR chip, which mimics cell-surface FcμR, Fcμ–J in the aqueous solution can participate in a multivalent interaction with FcμR, resulting in higher avidity and thus a higher apparent binding affinity. In contrast, FcμR in the mobile phase can only bind to the Fcμ–J in a monovalent manner, leading to an apparent binding affinity that reflects the average binding affinity of all the individual binding sites.

**Figure 3.**
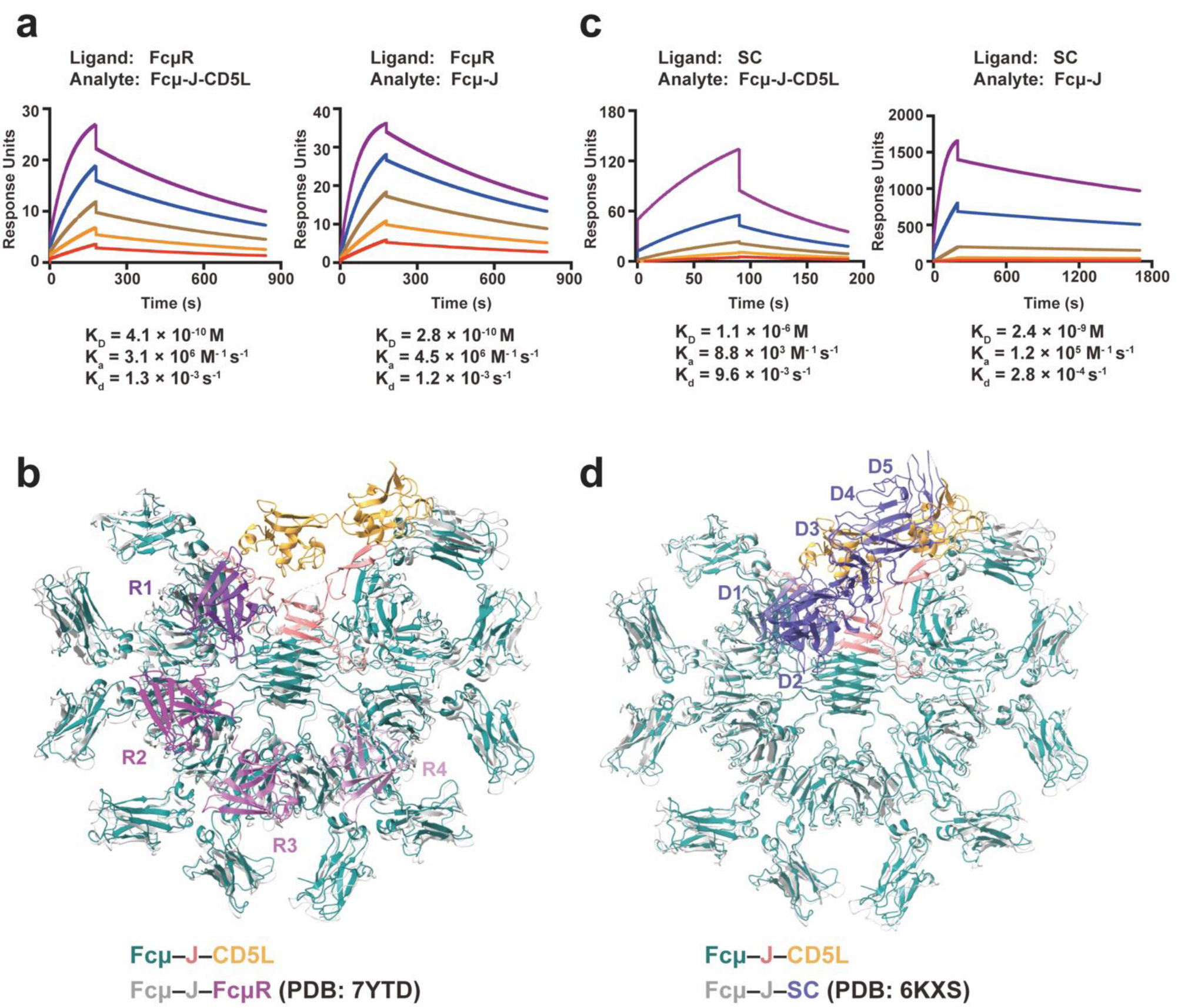
CD5L has no effect on the binding of FcμR but diminishes IgM binding to pIgR. **a.** SPR analyses of the interactions between immobilized FcμR and Fcμ–J or Fcμ–J–CD5L. **b.** Structural analyses suggest that CD5L would not interfere with the IgM–FcμR interaction. Fcμ–J and FcμR of the human Fcμ–J– FcμR complex (PDB: 7YTD) are shown in grey and purple, respectively. **c.** SPR analyses of the interactions between immobilized pIgR and Fcμ–J or Fcμ–J–CD5L. **d.** Structural analyses suggest that CD5L impede the IgM–pIgR interaction. Fcμ–J and pIgR of the human Fcμ–J–pIgR complex (PDB: 6KXS) are shown in grey and blue, respectively.

We were unable to measure the binding between IgM and FcαμR, the Fc receptor for both IgM and IgA^38^, due to challenges in producing well-behaved FcαμR protein. We then evaluated how CD5L influences the binding between IgM and the mucosal transport receptor, pIgR. Our results demonstrated that in the presence of CD5L, Fcμ–J associates with immobilized SC approximately 500-fold more weakly (Fig. 3c). Similar findings were observed when SC was used in the mobile phase (Extended Data Fig. 4b). It is noteworthy that the measured binding affinities between pIgR and Fcμ–J were consistent regardless of the stationary molecule, as pIgR binds to a single site in Fcμ–J. A structural comparison between Fcμ–J–CD5L and Fcμ–J–SC revealed that CD5L could substantially collide with the D3–D5 domains of SC (Fig. 3d), providing a rationale for the marked decrease in the interaction between Fcμ–J and SC in the presence of CD5L.

### CD5L does not affect IgM-mediated complement activation

IgM is capable of recruiting the C1q complex and initiating the classical complement pathway. The C1q binding site is situated in the Cμ3 domain of Fcμ, and it includes residues 432–436 in the FG loop^39^, which does not participate in interacting with CD5L. To access the effect of CD5L on IgM-mediated complement activation, we generated a recombinant anti-CD20 IgM molecule following previously established methods^40^. Subsequently, we conducted complement-dependent cytotoxicity experiments using OCI-Ly10 cells, which naturally express CD20. The results revealed that in the presence of human serum complement, the anti-CD20 IgM effectively induced the lysis of OCI-Ly10 cells (Fig. 4). Preincubation of this IgM with increasing amounts of CD5L did not show any significant effect on complement activation. In contrast, VAR2CSA, a malaria protein known for its potent inhibition of IgM-mediated complement cytotoxicity in this context^40^, effectively suppressed IgM-triggered cell killing. These findings suggest that CD5L does not exert a substantial influence on IgM-mediated complement activation.

**Figure 4.**
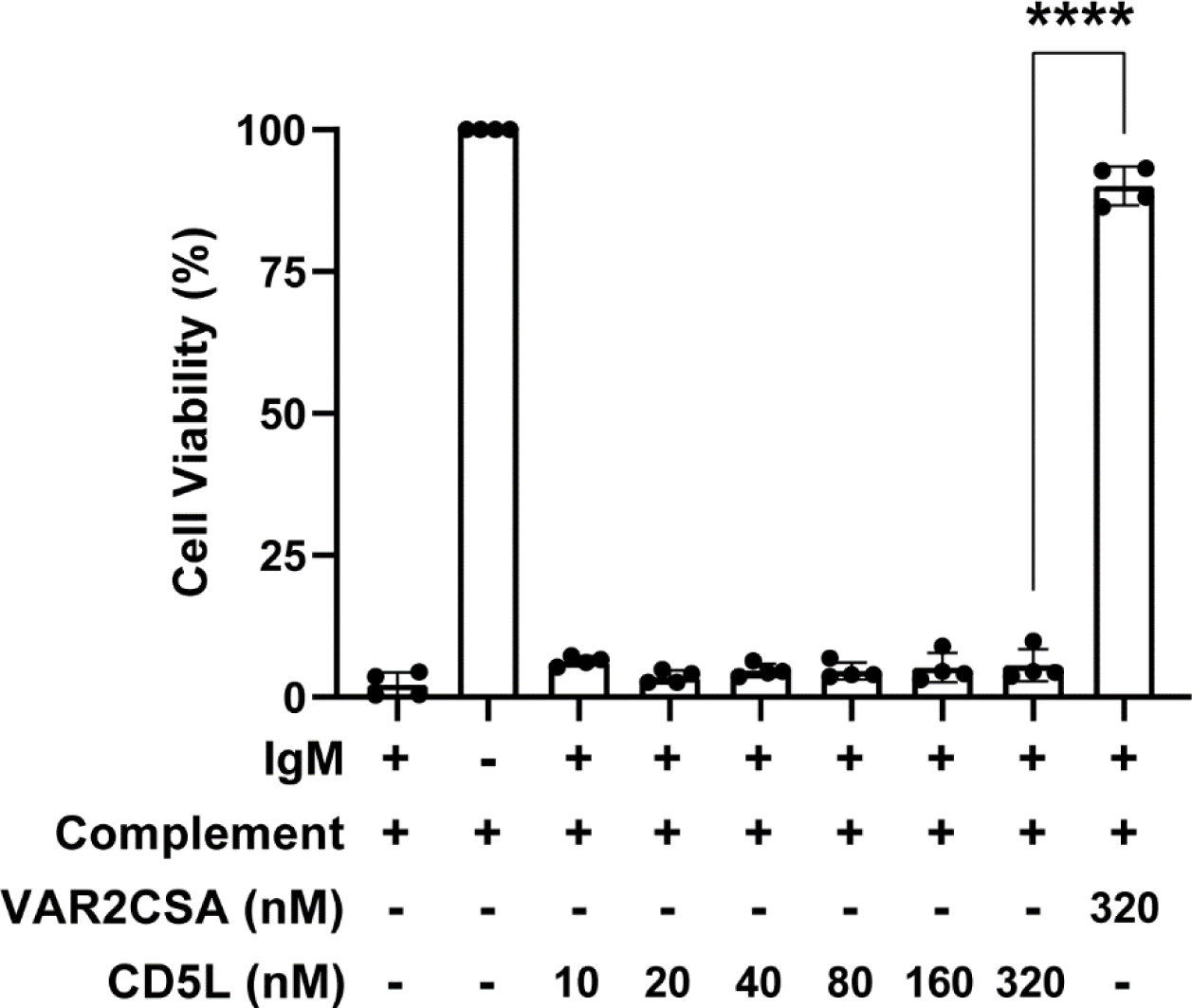
CD5L does not affect IgM-mediated complement activation. IgM-mediated complement activation was assessed using OCI-Ly10 cells, which naturally express CD20. Serially diluted CD5L (ranging from 320 nM to 10 nM) or the malaria protein VAR2CSA was incubated with anti-CD20 IgM (at 6 nM) for 3 hours. Subsequently, the protein mixtures were combined with normal human serum complement and applied to the OCI-Ly10 cultures. Cell viability was determined by measuring ATP levels using the CellTiter-Glo reagent. The experiment was conducted four times, and statistical analysis was performed using a two-tailed, unpaired Student’s t-test in GraphPad Prism. * indicates statistical significance at p ≤ 0.0001.

## Discussion

In this study, we have elucidated the molecular mechanism of the IgM–CD5L interaction. Our findings indicate that CD5L primarily engages with the J chain in a Ca^2+^-dependent manner, and the formation of a disulfide bond between CD5L and Fcμ further stabilizes the IgM–CD5L complex. Moreover, we have demonstrated that CD5L reduces IgM binding to pIgR but does not affect the binding of FcμR or complement activation. Collectively, these results corroborate recent findings by Oskam et al. and support the notion that CD5L is an integral component of circulatory IgM^8^.

Despite these advancements, the precise function of CD5L remains elusive. When bound to IgM, CD5L is shielded from renal elimination but appears to be functionally inactive. Upon disruption of immune homeostasis, CD5L dissociates from IgM and plays a role in immune regulation. Our results suggest that CD5L dissociation would likely require conditions that promote the disruption of the disulfide bond, as well as a decrease in Ca^2+^ concentration. Deciphering the molecular events that trigger CD5L release from IgM remains a critical area for future research. Furthermore, the existence of specific receptors for free CD5L is yet to be investigated. In any event, the fact that CD5L serves as a subunit of circulatory IgM alters our understanding of IgM and warrants immediate attention in related fields.

Finally, our investigation into the molecular basis of IgM–CD5L interaction has revealed that the J chain serves as a critical component, essentially functioning as an adaptor that enables the specific recognition of systemic IgA by FcRL4. The significance of the J chain was first reported by Marian Koshland in 1970, and in her 1985 review^41^, she highlighted that “the existence of a third immunoglobulin polypeptide, the J chain, is often overlooked.” Recent structural information obtained by our research and others has clearly demonstrated the essential role of the J chain in the polymerization of IgM and IgA, as well as its involvement in the recognition of IgM and IgA by pIgR. In this study and the accompanying manuscript, we have further revealed that the J chain also plays a crucial role in mediating the interaction between the IgM pentamer and CD5L, as well as between systemic IgA and FcRL4. It is evident that the J chain confers unique features to polymeric IgM and IgA, enabling them to be specifically recognized by distinct molecular partners to fulfill their functions. This emphasizes the pivotal role of the J chain in the immune system and highlights its significance in the specific recognition and function of polymeric immunoglobulins.

## Materials and Methods

### Protein expression and purification

The DNA fragment encoding the residues 20–347 of human CD5L (Uniprot O43866) was cloned into a modified pcDNA vector with a N-terminal IL-2 signal peptide followed by an 8-residue His tag. The plasmid expressing CD5L was transfected into the HEK293F cells by polyethylenimine (Polysciences), and the cells were cultured in SMM 293T-I medium (Sino Biological Inc.) at 37 °C, with 5% CO_2_ and 55% humidity. Four days following transfection, the conditioned media were collected by centrifugation, then concentrated via a Hydrosart Ultrafilter (Sartorius), and exchanged into the binding buffer (25 mM Tris-HCl, pH 7.4, 150 mM NaCl). The recombinant protein was isolated by Ni-NTA (Smart-Lifesciences) affinity purification and eluted with the binding buffer supplemented with 500 mM imidazole. The sample was further purified using a Superdex 200 increase column (GE Healthcare) and eluted using the binding buffer. Mouse CD5L (residues 22–352, Uniprot Q9QWK4) and cat CD5L (3-SRCR isoform, residues 20–355, Uniprot A0A1E1GEY0) were produced similarly.

Human Fcμ–J complex was expressed and purified as previously described^30^. Cat Fcμ (residues 94–441, NCBI BAA32231.1)–J (residues 1–158, Uniprot M3XBL5) and mouse Fcμ (residues 104–454, Uniprot P01872)–J (residues 1–159, Uniprot P01592) complexes were prepared in similar manners. To obtain the human Fcμ–J–CD5L tripartite complex, purified CD5L was mixed with Fcμ–J complex at a 3:1 molar ratio and the mixture was incubated at 4 ℃ overnight. The target complex was then further purified on a Superose 6 increase column and eluted using the binding buffer supplemented with 5 mM CaCl_2_.

### Surface plasmon resonance

To verify the interaction of CD5L and Fcμ–J, the surface plasmon resonance experiments were performed on the Biacore T200 (GE Healthcare). Fcμ–J was immobilized on a CM5 chip (GE Healthcare) at pH 4.5 using the amine coupling procedure at a 10 μL/min flow rate. CD5L was injected over the flow cell at a range of five concentrations prepared by serial twofold dilutions at a flow rate of 30 μL/min using a single-cycle kinetics program. The SPR assays were performed at 25 °C using the following running buffer: 10 mM HEPES, pH 7.4, 150 mM NaCl, 3 mM EDTA, 0.05% Tween-20, with or without 8 mM CaCl_2_.

To investigate the effect of the interaction of Fcμ–J–CD5L with IgM receptors, SC and the D1 domain of FcμR was first immobilized on the chip: SC was biotinylated at the C-terminal and immobilized on the SA chip (GE Healthcare), whereas FcμR-D1 was immobilized on the CM5 chip to a level of resonance units similar to pIgR. Fcμ–J–CD5L and Fcμ–J was then injected over the flow cell respectively at a range of five concentrations prepared by serial twofold dilutions at a 30 μL/min flow rate using a high-performance program. The experiments were also carried out conducted in the reverse manner, with Fcμ–J–CD5L and Fcμ–J immobilized on the CM5 chip, respectively. Binding studies were then performed by passing twofold serial dilutions of purified pIgR or FcμR over the chip. This assays were performed with a running buffer (10 mM HEPES, pH 7.4, 150 mM NaCl, 3 mM EDTA, 8 mM CaCl_2_, 0.05% Tween-20) at 25 ℃.

All data were fitted to a 1:1 binding model using Biacore Evaluation Software. Each SPR experiment was repeated at least three times.

### Cryo-EM sample preparation and data collection

Prior to cryo-EM sample preparation, the gel filtration peak fractions containing Fcμ–J–CD5L tripartite complex was collected and concentrated to 0.25 mg/mL, then cross-linked with 0.05% glutaraldehyde (Sigma) at 20℃ for 5 min. For grid preparation, 4 μL of Fcμ–J–CD5L was applied to glow-discharged holy-carbon gold grids (Quantifoil, R1.2/1.3, 300 mesh), blotted with filter paper in 100% humidity at 4 °C, and put into the liquid ethane with a Vitrobot Mark IV (FEI). A Talos Arctica microscope equipped with Ceta camera (FEI) was used to screen grids. Data collection was performed on a Titan Krios electron microscope (FEI) operated at 300 kV with the Gatan Imaging Filter (GIF) energy-filtering slit width set at 20 eV. Movie stacks were recorded on a K3 Summit direct electron detector (Gatan) in super-resolution mode at a magnification of 105,000× using the EPU software (E Pluribus Unum, Thermo Scientific), which corresponded to a pixel size of 0.83 Å per pixel. The dose rate was 16 e^–^/pixel/s. The defocus range was set from –1.0 to –1.5 μm. The micrographs were recorded for 2.56 s and subdivided into 32 frames with a total electron exposure of 59.5 electrons per Å^2^. Statistics for data collection are summarized in Extended Data Table 1.

### Imaging processing, model building and structure refinement

For 3D reconstruction of the Fcμ–J–CD5L complex, a total of 6,962 movie stacks were recorded. All image processing was performed with cryoSPARC^42^. Raw movie frames were motion-corrected and dose-weighted using Patch motion correction (multi) module. The contrast transfer function (CTF) parameters of each summed image were estimated by Patch CTF estimation (multi) module. The images were screened manually to remove low-quality ones with the Manually Curate Exposures module. A set of 979,111 particles were picked without reference by Blob picker module and subjected to reference-free 2D classification to generate templates for template particle picking. Subsequently, a total of 894,255 particles were template-picked, and extracted particles were subjected to several rounds of 2D classification to remove noise and fuzzy particles. 769,364 particles were kept for initial model generation and 3D classification, using Ab-Initio Reconstruction and Heterogeneous Refinement, respectively. The particles in qualified groups (381,470 particles) were combined and subjected to Homogeneous Refinement, resulting in a map with a 3.39 Å overall resolution, according to the Fourier shell correlation (FSC) = 0.143 criterion. Local Refinement was performed with a soft mask which encompasses the CD5L-Fcμ/J interface, yielding a map at the resolution of 3.41 Å with a clearer visualization of the interface. The mask was created using UCSF chimera^43^. The local resolution map was analyzed by Local Resolution Estimation module of cryoSPARC and displayed using UCSF ChimeraX.

The initial model of CD5L was predicted by Alphafold^44^, as well as the structures of Fcμ from the Fcμ– J–SC complex (PDB ID: 6KXS) and J chain from the SIgA complex (PDB ID: 6LX3), was docked into the cryo-EM map using UCSF chimeraX and then manually adjusted using Coot^45^. The SRCR1 domain of CD5L is absent in the structure because of displaying poor densities, perhaps due to potential conformational flexibility. Refinement was performed using the real-space refinement in Phenix^46^. Figures were prepared with UCSF ChimeraX.

### StrepTactin pull-down assay

CD5L mutations were introduced through PCR-based mutagenesis. Afterwards, wild type and mutant CD5L proteins were purified with the Ni-NTA affinity method. For the pull-down assays, CD5L was first incubated with purified Fcμ–J complex on ice for 1 h. The mixture was then incubated with the StrepTactin beads (Smart Lifesciences) and rotated at 4 °C, in the binding buffer containing 25 mM Tris-HCl, pH 7.4, 150 mM NaCl, 5 mM CaCl_2_ or 5 mM EDTA. A twin-strep tag is present on Fcμ. After 1 h of incubation, the beads were washed three times with the binding buffer, and the bound proteins were eluted with the binding buffer supplemented with 10 mM desthiobiotin (IBA Lifesciences). The samples were analyzed by SDS-PAGE and detected by Coomassie staining.

### Complement-dependent cytotoxicity assay

Anti-CD20 IgM was produced as described previously^40^. OCI-Ly10 cells, which express CD20, were used for the complement-dependent cytotoxicity assay. Serially diluted CD5L were incubated with anti-CD20 IgM for 3 hours in 50 μL of RPMI-1640. Then the proteins were mixed with equal amounts of normal human serum complement (1:12.5, Quidel) and OCI-Ly10 cultures (20,000 cells) before being put into a 96-microwell plate. 50 μL of CellTiter-Glo reagent (Promega, G7572) was added to each well and incubated for 10 min at room temperature after 6 h of incubation at 37 °C. A Cytation 5 cell imaging multi-mode reader (BioTek) was used to quantify luminescence. The experiment was repeated four times and the statistical were analyzed by two-tailed, unpaired Student’s t-test in GraphPad Prism.

## Data and materials availability

Cryo-EM density maps of Fcμ–J–CD5L have been deposited in the Electron Microscopy Data Bank with accession codes EMD-37936 (overall) and EMD-37937 (local). Structural coordinates have been deposited in the Protein Data Bank with the accession codes 8WYR (overall) and 8WYS (local).

## Acknowledgments

We are grateful to the Core Facilities at the School of Life Sciences, Peking University for their assistance with negative-staining EM; the Cryo-EM Platform of Peking University for their support with data collection; and the High-performance Computing Platform of Peking University for their aid with computation. We also acknowledge the National Center for Protein Sciences at Peking University for their help with Biacore and BioTek facilities. This work received support from the National Natural Science Foundation of China (32325018) and the Qidong-SLS Innovation Fund to J.X., as well as from Changping Laboratory.

## Author Contributions

Y.W. and C.S. contributed equally to this work. Y.W. and C.S. performed protein expression, purification, cryo-EM sample screening and data collection. Y.W. carried out the cryo-EM data processing, model building and structure refinement. C.S. performed the SPR analyses and the complement-dependent cytotoxicity assay with assistance from C.J. Y.W. and C.S. conducted the pull-down experiments. J.X. conceived and supervised the project and wrote the manuscript, with inputs from all authors.

---

**Extended Data Fig. 1.**
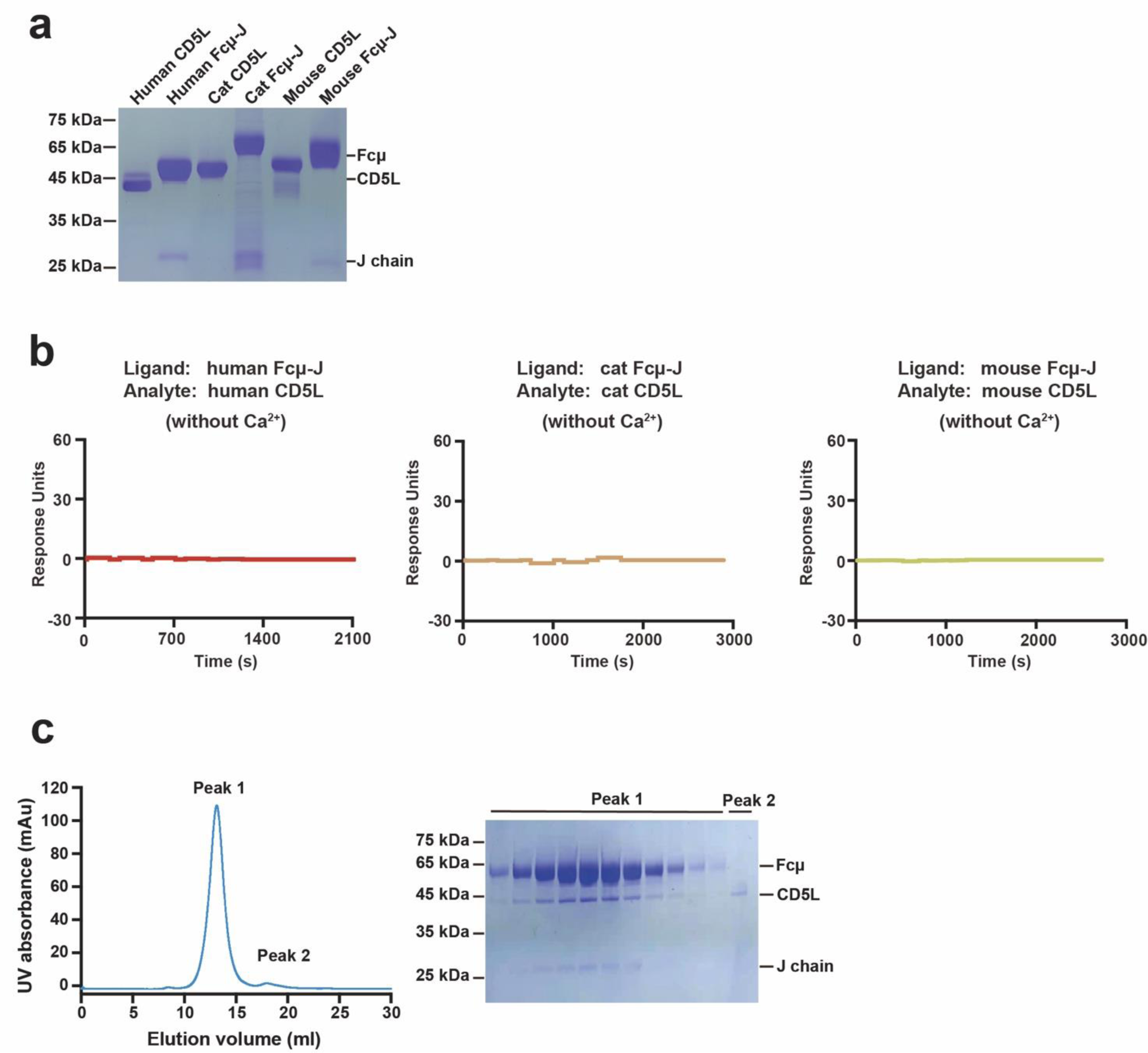
Purification of the CD5L and Fcμ–J. **a.** SDS–PAGE analyses of the CD5L and Fcμ–J form human, mouse and cat. **b.** No significant interaction between CD5L and Fcμ–J is detected in the presence of EDTA. **c.** Size-exclusion chromatography of the Fcμ–J–CD5L complex on a Superose 6 Increase column and SDS– PAGE analyses.

**Extended Data Fig. 2.**
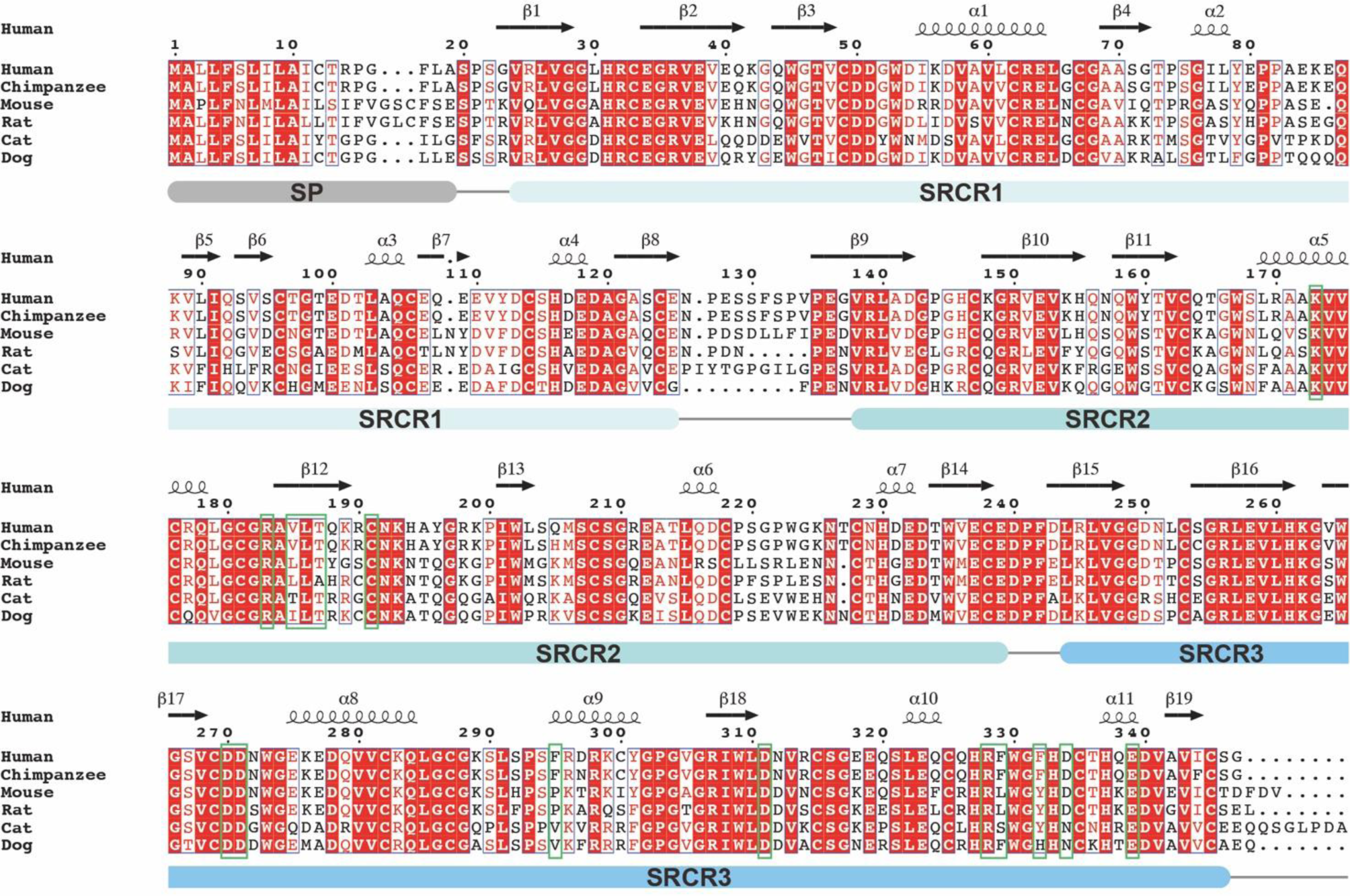
Sequence alignment of CD5L. Sequence alignment of CD5L from human and other animals. Residues in human CD5L that are involved in binding to Fcμ–J are highlighted in green boxes. Signal peptide (SP) is shown in grey. Three SRCR domains are shown in light cyan, light blue and blue, respectively.

**Extended Data Fig. 3.**
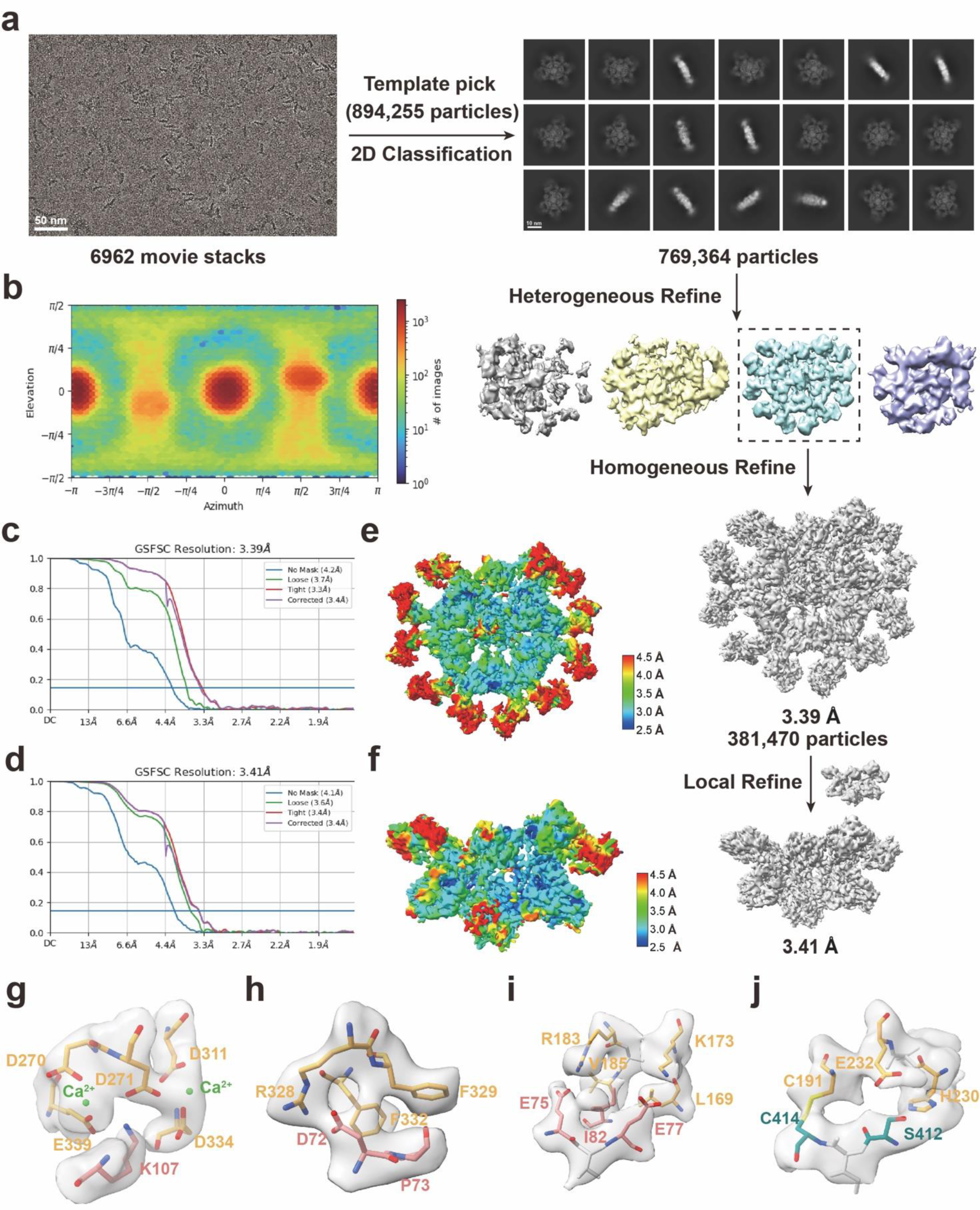
Cryo-EM 3D reconstruction of the Fcμ–J–CD5L complex. **a.** Flowchart of cryo-EM data processing. **b.** Angular particle distribution heat map. **c-d.** Gold-standard Fourier shell correlation (GSFSC) curves of the global and local Fcμ–J–CD5L complex, respectively. **e-f.** Resolution estimations for the final maps of the global and local Fcμ–J–CD5L complex, respectively. **g.** The DDE/D motif of CD5L-SRCR3 interacts with J chain. **h-i.** The SRCR2–SRCR3 junction of CD5L interacts with the β5–β6 hairpin of J chain. **j.** CD5L-SRCR2 interacts with Fcμ5B.

**Extended Data Fig. 4.**
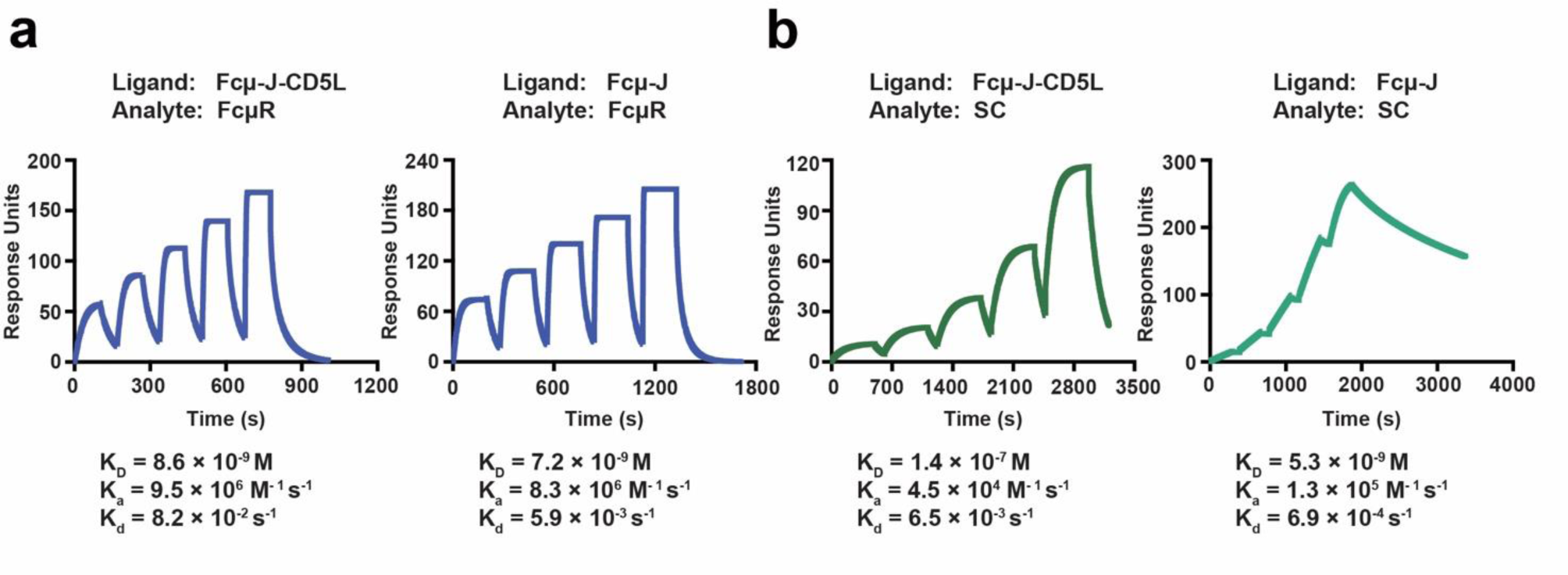
SPR analyses using immobilized Fcμ–J or Fcμ–J–CD5L. **a.** SPR analyses of the interactions between immobilized Fcμ–J or Fcμ–J–CD5L and FcμR. **b.** SPR analyses of the interactions between immobilized Fcμ–J or Fcμ–J–CD5L and pIgR.

**Extended Data Table 1.**
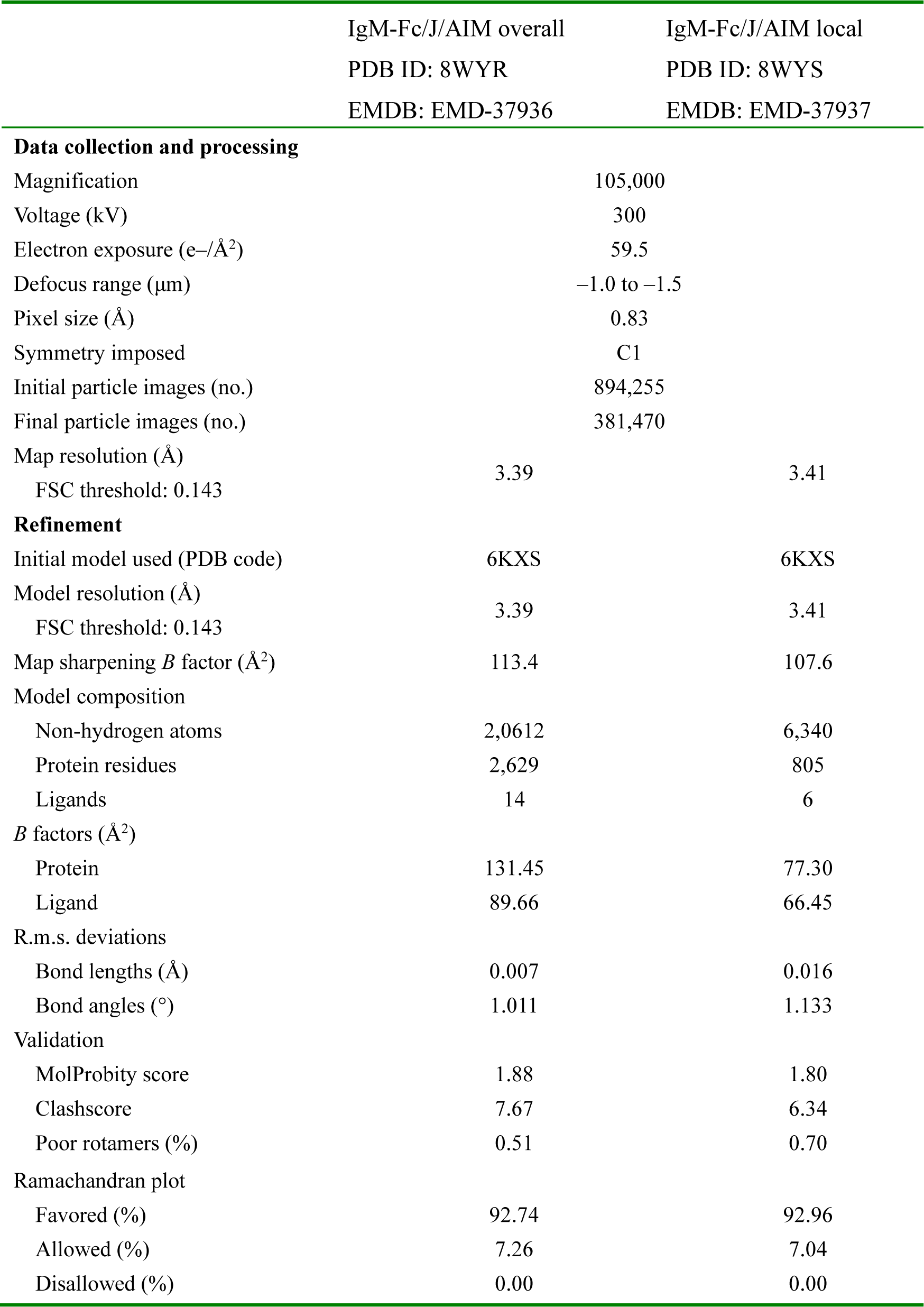
Cryo-EM data collection, refinement and validation statistics.

